# Lifetime development changes in rats tracked by urinary proteome

**DOI:** 10.1101/2022.09.21.508856

**Authors:** Xuanzhen Pan, Yongtao Liu, Yijin Bao, Youhe Gao

## Abstract

The existing researches mainly focus on the embryonic stage and a short time after that. However, there was little research about the whole life of an individual from childhood to aging and death. For the first time, we used non-invasive urinary proteome technology to track the changes of several important development timepoints in a batch of rats, covering the 10 timepoints from childhood, adolescence, young, middle adulthood, and to old age death. As previous studies on puberty found, sexual or reproductive maturation, mature spermatozoa in seminiferous tubules(first seen), gonadal hormones, decline of estradiol(E), brain growth and central nervous system myelination, and our differential protein enrichment pathways also included reproductive system development, tube development, response to hormone, response to estradiol, brain development, neuron development.

Like previous studies found in young adulthood, musculoskeletal maturity, peak bone mass, development of the immune system, and growth and physical development, and our differential protein enrichment pathway also included skeletal system development, bone regeneration, system development, immune system process, myeloid leukocyte differentiation, growth, and developmental growth. However, at all timepoints in the whole life, there were still many biological pathways of the differential urinary protein enrichment involving multiple organs, tissues, systems, etc., which had not been mentioned by existing studies. This study provided comprehensive and detailed changes in rats’ lifetime development through urinary proteome, supplementing the blank of development research. But also provided a new idea for monitoring the human health change condition and the possibly occurring aging diseases by using the urinary proteome.

**Summary:** As far as we know, this is the first time that the urinary proteome was used to track the changes of several important developmental timepoints in the lifetime of rats.

## Introduction

Urine is considered one of the most valuable biofluids for the discovery of disease biomarkers because urine collection is noninvasive and easy. More importantly, unlike blood, urine is not subjected to homeostatic control, and it accumulates small, sensitive, and early changes associated with systemic changes, some of which may be used as biomarkers1 (Gao, 2013). Urine proteomics has already been applied to various clinical studies (Rodríguez-Suárez et al., 2014; Zou and Sun, 2015), including studies on lung cancer (Chen et al., 2015; Wang et al., 2017; Zhang et al., 2015; Zhang et al., 2018), breast cancer (Beretov et al., 2015; Gajbhiye et al., 2016), bladder cancer (Duriez et al., 2017; Lei et al., 2013; Santoni et al., 2018), gastric cancer (Shimura et al., 2020), genitourinary cancer (Chen et al., 2019), and knee osteoarthritis (Xiao et al., 2019). Moreover, urine-filtered plasma proteins originate from distal organs, including the brain, etc. not only the kidney (An and Gao, 2015; Decramer et al., 2008; Winter et al., 2020).

Most modern developmental biology research has focused on individuals during pregnancy and a short time before reaching adulthood and generally focuses on a certain organ or system (Gong et al., 2020; Li et al., 2018). There were also some studies on elderly individuals (Stanley and Shetty, 2004), but they only involved a certain organ, and individuals of different ages were different, so there might be a lack of comparability. This study is the first time that urinary proteome was used to track the whole development of a batch of rats from childhood to old age. Urinary proteome can also reflect the overall changes of the body.

## Results

### Overview of all rat cohorts for urinary proteome

All rat cohort was assessed using label-free data□independent acquisition (DIA) LC-MS/MS quantification to characterize the urinary protein profile (Gillet et al., 2012; Ludwig et al., 2018) (**Fig. 1a**). To map proteome changes between individuals with different timepoints, we analyzed 144 urine samples (including 126 individual samples and 18 mix samples) from rat cohorts. To maximize proteome depth, a spectral library was generated by merging three sub□libraries: (i) a library constructed by data□dependent acquisition (DDA) consisting of 10 fractions of pooled urine samples; (ii) a DDA library converted from consisting of 10 fractions of urine samples by PD; and (iii) a library generated from the DIA analysis of all analyzed samples. A total of 2960 protein groups were identified in all samples of the rat cohort.

**Fig. 1.**
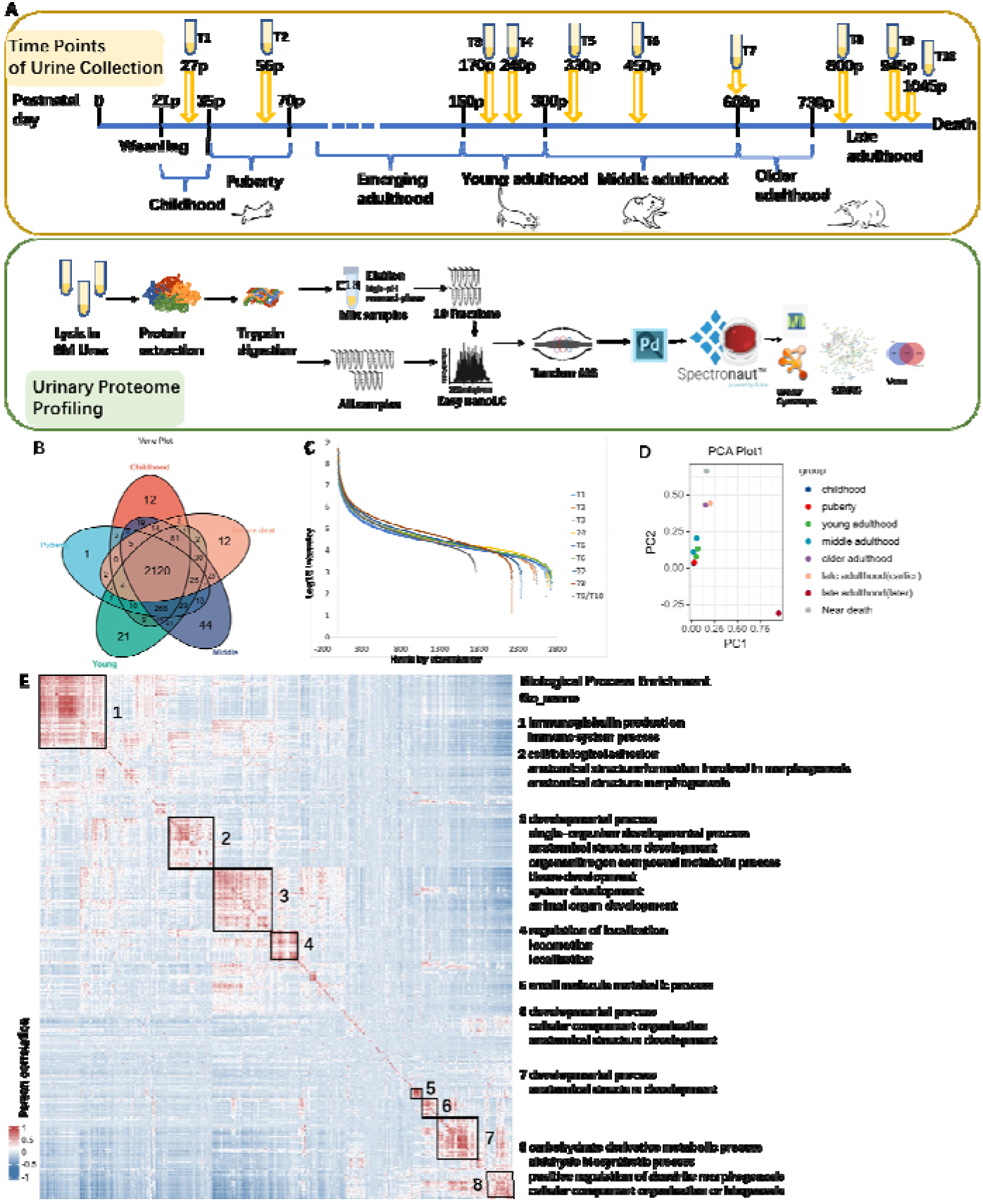
Urinary proteome profiling of the developing rats. A. Illustration of rats’ development stages and 15 sampling timepoints. Experimental procedure and data analysis workflow below. B. The overlap of proteins that were identified in five stages, including childhood, puberty, young, middle and near death. Common proteins identified among all stages and stages exclusive proteins were present. C. Dynamic ranges of the rat urinary proteome measured at 10 timepoints. Proteins identified in the all stage cohorts were ranked according to their MS signals, which covered more than five orders of magnitude. D. Principal component analysis of the temporal proteome data, including 10 timepoints. Time points before middle age get together. E. Global correlation map of proteins generated by clustering the Pearson correlation coefficients of all pairwise protein comparisons.

All rat samples were collected from ten timepoints. Among them, the first timepoint(T1) was in childhood, the second timepoint(T2) was in adolescence, the third and fourth timepoints(T3, T4) were in young adulthood, the fifth and sixth timepoints(T5, T6) were in middle adulthood, the seventh timepoint(T7) began reproductive decline, the eighth timepoint(T8) was in later life, only three rats were left by the ninth timepoint(T9), and the tenth timepoint(T10) was taken two days before the death of the last rat(detailed information in **Fig. 1A**). While some proteins are unique in each period, but 2120 proteins were identified in all periods (**Fig. 1B**). The quantified protein intensities spanned five orders of magnitude in all timepoints cohorts and the top ten most abundant proteins contributed about half to the total urinary proteome signal (**Fig. 1C**). To examine the data quality of all timepoint samples, principal component analysis (PCA) was conducted (**Fig. 1D**), in which before middle adulthood, the timepoints gather together, while the timepoints between later life and before death were far apart. The global correlation map contained pairwise relations of all urinary proteins across 126 samples from all timepoints cohorts. Unsupervised hierarchical clustering of the pairwise Pearson correlation coefficients revealed some small clusters of co-regulated proteins (**Fig. 1E**). These clusters were chiefly enriched for proteins with the Gene ontology (GO)-terms as well as other significant terms. The biological processes included some immune system processes, development, morphogenesis, localization, and metabolic process.

### The same features between existing researches and urinary development research

Unsupervised hierarchical clustering of Pearson correlation coefficients reveals that most timepoints are arranged in sequence, which is consistent with the development or aging with time (**Fig. 2A**). To achieve higher confidence, differential proteins identified in at least two of the seven biological replicates in at least one-time point were used for further statistical analyses. Differential proteins also met the conditions *p*-value <0.05 and fold change >2 or <0.5. The differential proteins were compared between the two timepoints, and all the compared time points were shown in **Fig. 2A**. The cross-over relationship of differential proteins in the five stages was shown in **Fig. 2A**. There were 36 differential proteins in common in the five stages, and many proteins were only produced in certain stages. Four rats died shortly after 800 days, two died shortly after 945 days, and the remaining rat lived for 147 days. Each rat’s late development and aging process were different, which reflected certain individual differences.

**Fig. 2.**
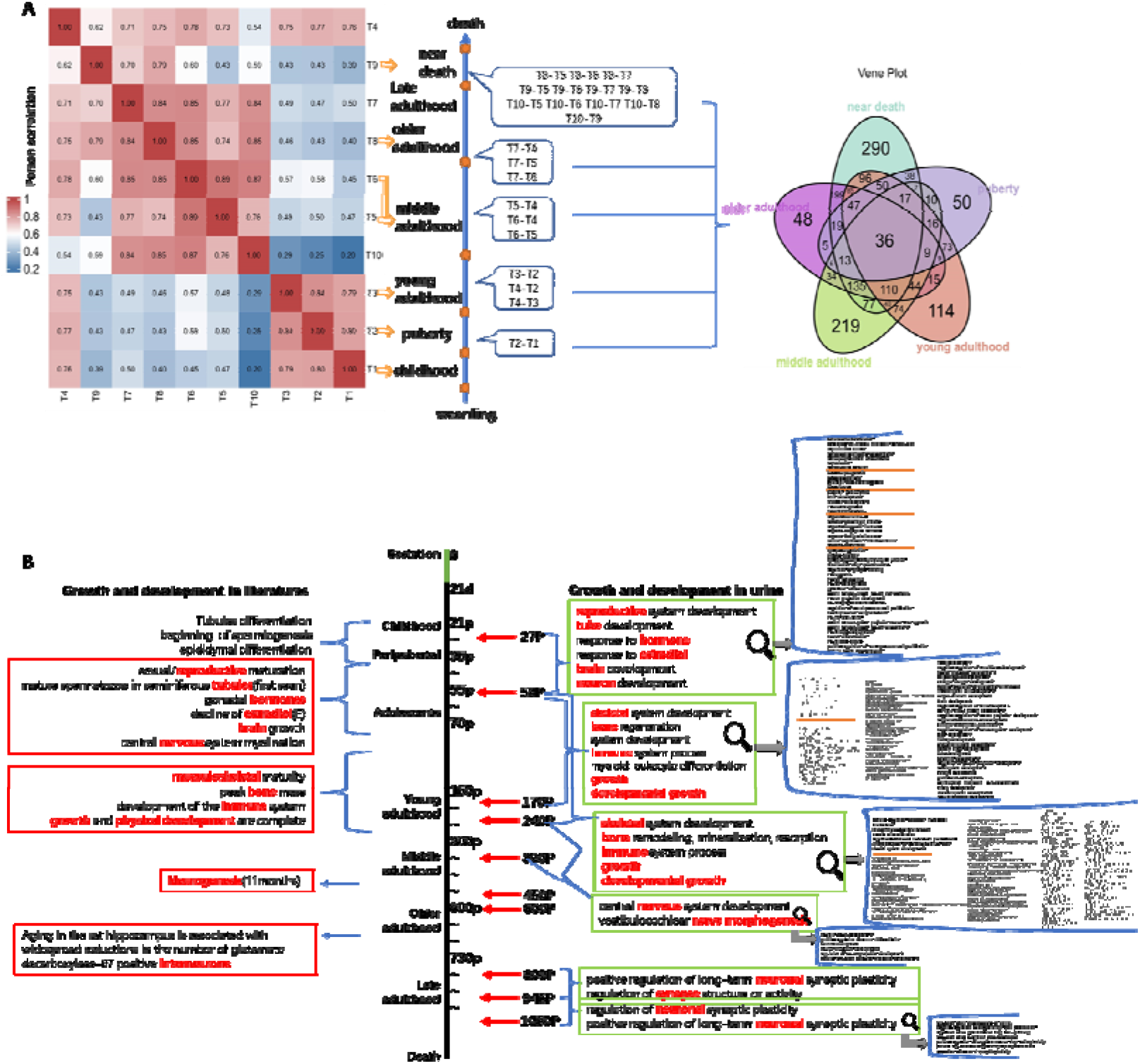
Comparison between existing researches and urinary development research. A. Hierarchical clustering analyses of temporal proteome data separated the rat development into six phases. And the process of obtaining and analyzing differential proteins. The middle shows the comparison of all time points. The overlap of differential proteins that were identified in five stages (fold change > 2 or < 0.5, p-value <0.05). B. The Panel showed comparisons of previous studies involving various stages of development and urinary development research for mutual validation. The displayed biological pathways met the requirements p-value adjusted < 0.01.

The differential proteins from samples comparison between the two timepoints were analyzed separately for enrichment gene ontology (GO) analysis processed by OmicsBean (Binns et al., 2009). Compared with previous research results, we found that all the biological pathways enriched by urine differential proteins can find corresponding pathways, which could be mutually verified, as shown in **Fig. 2B**. As mentioned in previous studies, the peripubertal period occurs from the onset of puberty, where circulating gonadal hormones start to rise, leading to sexual/reproductive maturation (Bell, 2018; Picut et al., 2018); puberty is the developmental stage in which sexual development is completed, and reproductive capacity or fertility is achieved (Ojeda et al., 1980; Vidal, 2017); the onset of puberty in male rats is when mature spermatozoa are first seen in seminiferous tubules (Picut et al., 2018); estradiol increased in peripubertal period in rats, maintaining expression in adolescence (Bell, 2018); significant brain growth is ongoing in the rat until nine weeks of age and central nervous system myelination in limbic structures is not complete until six weeks of age (Bandeira et al., 2009; Downes and Mullins, 2014; Jackson et al., 2017). Accordingly, the differential proteins obtained on days 56 and 27 were compared and enriched to a similar biological pathway: reproductive system development; tube development; response to hormone; response to estradiol; brain development; neuron development (*p*-value adjusted < 0.01, only a few major pathways were listed here, and more detailed information was presented in **Supplementary Table 1**).

Research on rats that enter adulthood after puberty found that: adulthood in rats is determined according to musculoskeletal maturity (Quinn, 2005), and adult life is after growth and physical development are complete (Roe et al., 1995); unlike humans, bone growth never completely stops in rats (SIMSON and GOLD, 1982); peak bone mass is not reached until around 26 weeks of age in rodents (Brodt et al., 1999; Halloran et al., 2002; Somerville et al., 2004); the development of the immune system is defined by changes in thymus size and cellular content over early development as well as key immunological markers and T and B-lymphocyte production increases over the first 26 weeks of life (Advances in Immunology Volume 31, 1981; Holsapple et al., 2003; Kay et al., 1979; Kincade, 1981). Similarly, the biological pathways enriched by the differential proteins obtained by comparing young adulthood with adolescence also contained these processes: skeletal system development, bone regeneration, system development, immune system process, myeloid leukocyte differentiation, growth, developmental growth, bone remodeling, mineralization, and resorption.

Some studies also involved the research of neurogenesis or interneurons in middle-aged and old-aged rats (Merkley et al., 2014; Stanley and Shetty, 2004). We also found similar biological pathways, such as central nervous system development, vestibulocochlear nerve morphogenesis, regulation of neuronal synaptic plasticity, and positive regulation of long-term neuronal synaptic plasticity in the comparison between middle adulthood and young adulthood and between late adulthood and older adulthood.

We also conducted the literature search for significantly changed differential proteins and found proteins at various time points that were related to the development stages mentioned in the previous studies. The specific proteins and descriptions of development stages related to the literature are shown in **Supplementary Table 2**.

### Presentation of development stages in urinary proteome

Enrichment analysis was performed on the differential proteins (fold change > 2 or < 0.5, p-value <0.05) obtained by pairwise comparison at adjacent timepoints in each stage. As shown in **Table 1**, biological pathways are common to at least three stages (p-value adjusted < 0.01, PAS Zscore means the pathway activation strength Zscore, served as the activation profiles of the Signaling pathways based on the expression of individual genes). Among them, all timepoints had in common were tissue development and system development. Anatomical structure development/morphogenesis, blood vessel development/morphogenesis, cardiovascular system development, circulatory system development, response to hormone, epithelium development, animal organ development, angiogenesis, regulation of growth, urogenital system development, immunoglobulin production, response to steroid hormone, response to peptide hormone, cellular response to growth factor stimulus, epithelial cell differentiation, positive regulation of cell differentiation, positive regulation of cell proliferation, regulation of vasculature development also involved five stages (More detailed information for all timepoints was presented in **Supplementary Table 1**). And most of these were not mentioned in the existing researches.

**Table 1:**
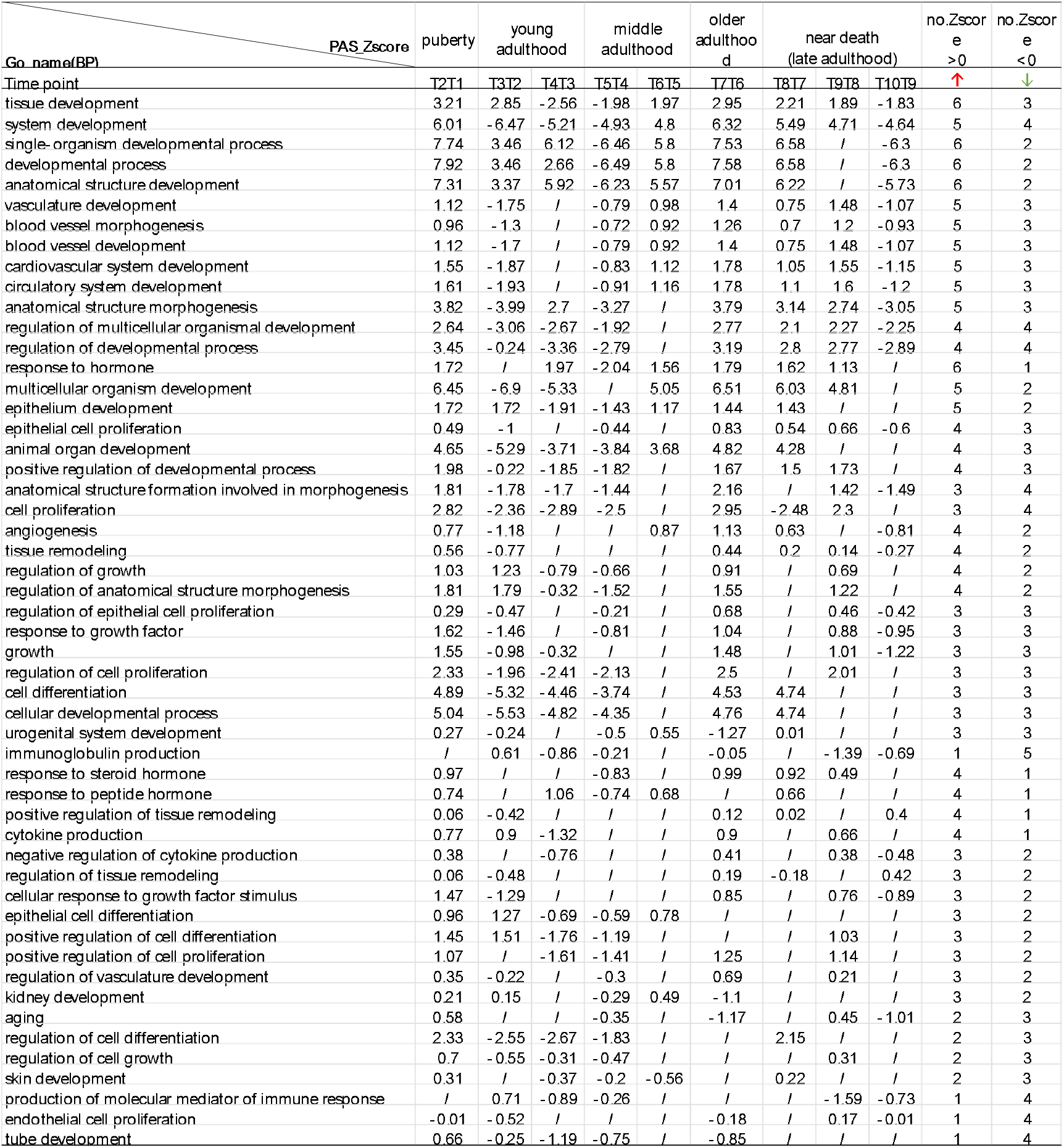
Enrichment analysis on the differential proteins in five stages.

We compared the obtained differential proteins at the 10 timepoints for enrichment analysis, respectively. The enriched biological pathways are mostly involved in development, growth, morphogenesis, formation, remodeling, regeneration, differentiation, and others (*p*-value adjusted < 0.01). **Fig. 3** shows the biological pathways involved in most stages, and there were many pathways involving fewer stages, as shown in **Supplementary Table 3**. It was found that the biological pathways involved aging, and modulation of age-related behavioral decline. We also found that the pathways involved in the growth and development of many organs, tissues, or systems, such as the brain, neurons, bone, kidney, liver, gland, cardiovascular system, hormone, etc. All specific biological pathway information and Z-score of pathways were shown in **Supplementary Table 3**.

**Fig. 3.**
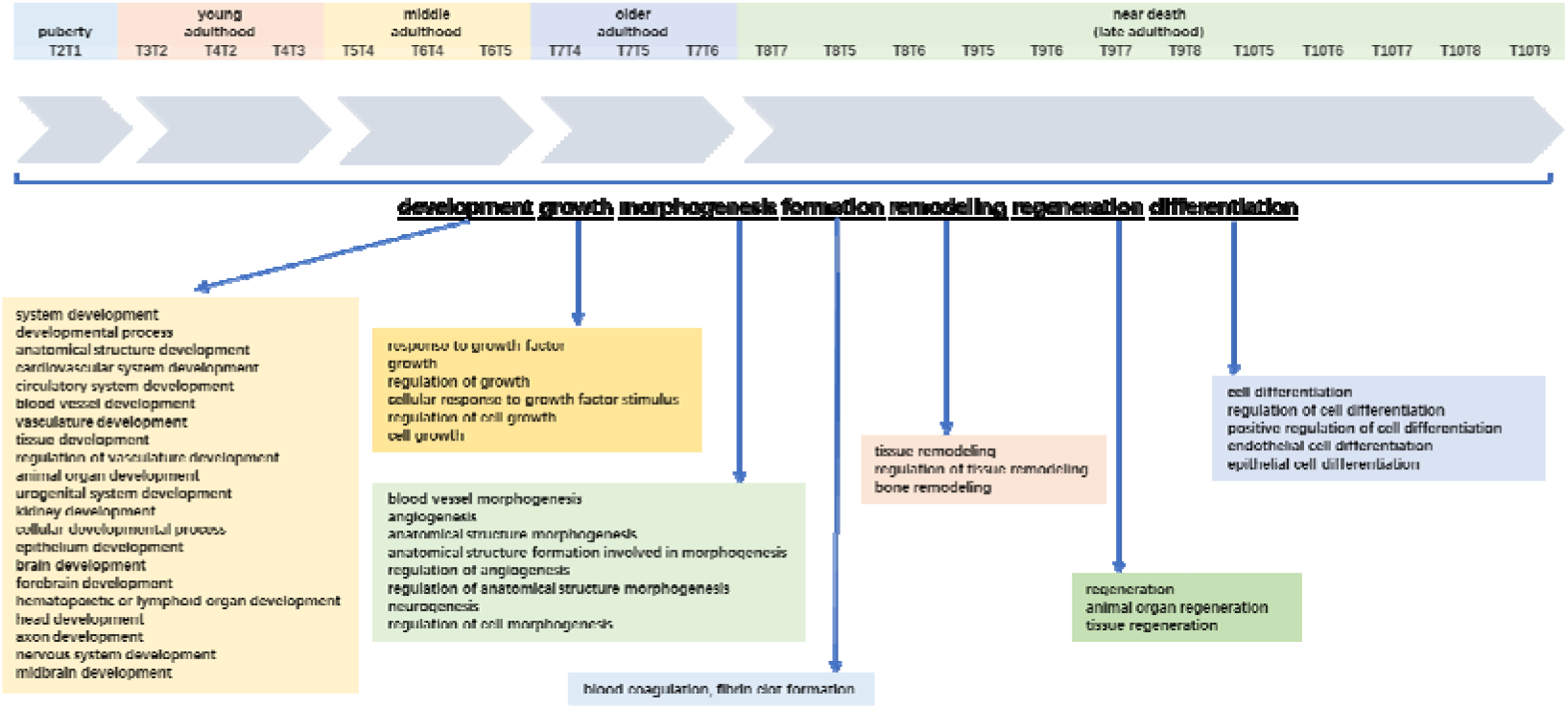
Overview of the biological pathway analysis of differential protein enrichment obtained by comparison at all timepoints. Biological pathways involved many organs, tissues or systems and spanned many biological processes such as development, growth, differentiation, remodeling, etc. Only some major biological processes covering five development stages were shown in this figure (p-value adjusted < 0.01), and the comprehensive biological processes were shown in the Supplementary Table 2.

## Discussion

In this study, we used the urinary proteome for the first time to monitor the growth and development of a batch of rats from childhood to death. The urinary proteome comprehensively reflected the growth and development of each stage, and almost covered all aspects of the body from organs to tissues to the system.

In comparison with previous studies, we found a number of body changes that were not mentioned in previous studies in many stages. We also found changes in some organs with age and aging-related pathways covering almost all timepoints. Urine is non-invasive and convenient. It is believed that this experiment can become a highlight in the field of growth and development, demonstrating for the first time that urine can show changes in body growth and development in all aspects, and providing ideas for monitoring body conditions of patients and aging research, such as clinical prognosis in the future.

## Materials and methods

### Acquisition of experimental samples and data

The rat cohort includes included 7 male rats of which the mother and father were born from the same brood of the same parents. We collected data from their major developmental periods including childhood(27 days), adolescence(56 days), youth(170 days, 240 days), adulthood(330 days, 450 days, 600 days) to old age death(800 days, 945days, 1050 days) (Andreollo et al., 2012; Ghasemi et al., 2021; Sengupta, 2013). All rats were bred from birth to this the indicated day, with the same fodders, and they lived in the same environment. The animal experiments were approved by the Ethics Review Committee of the Institute of College of Life Science, Beijing Normal University, China. Male rats’ parents were purchased from Beijing Charles River Laboratory. The rats were acclimated to the environment for one week before the experiment. All experimental animals were utilized following the “Guidelines for the Care and Use of Laboratory Animals” issued by the Beijing Office of Laboratory Animal Management (Animal Welfare Assurance Number: ACUC-A02-2015-004). These urine samples were collected in the same environment, and they were frozen in -80□fridge refrigerators, and then were processed together.

### Urine sample preparation for label-free analysis

After collection, the urine samples were centrifuged at 3000 ×g for 30 min at 4 °C and then stored at − 80 °C. For urinary protein extraction, the urine samples were first centrifuged at 12,000 ×g for 30 min at 4 °C. Then, 15 mL of urine from each sample was precipitated with three volumes of ethanol at − 20 °C overnight. The pellets were dissolved in lysis buffer (8 mol/L urea, 2 mol/L thiourea, 50 mmol/L Tris, and 25 mmol/L dithiothreitol). Finally, the supernatants were quantified by the Bradford assay.

A total of 100 μg of protein was digested with trypsin (Trypsin Gold, Mass Spec Grade, Promega, Fitchburg, WI, USA) using filter-aided sample preparation (FASP) methods (Wiśniewski et al., 2009). The protein in each sample was loaded into a 10-kDa filter device (Pall, Port Washington, NY, USA). After washing two times with urea buffer (UA, 8 mol/L urea, 0.1 mol/L Tris-HCl, pH 8.5) and 25 mmol/L NH4HCO3 solutions, the protein samples were reduced with 20 mmol/L dithiothreitol at 37 °C for 1 h and alkylated with 50 mmol/L iodoacetamide (IAA, Sigma) for 45 min in the dark. The samples were then washed with UA and NH4HCO3 and digested with trypsin (enzyme-to-protein ratio of 1:50) at 37 °C for 14 h. The digested peptides were desalted using Oasis HLB cartridges (Waters, Milford, MA, USA) and then dried by vacuum evaporation (Thermo Fisher Scientific, Bremen, Germany).

The digested peptides were dissolved in 0.1% formic acid and diluted to a concentration of 0.5 μg/μL. To generate the spectral library for DIA analysis, a pooled sample (1∼2 μg of each sample) was loaded onto an equilibrated, high-pH, reversed-phase fractionation spin column (84,868, Thermo Fisher Scientific). A step gradient of 8 increasing acetonitrile concentrations (5, 7.5, 10, 12.5, 15, 17.5, 20, and 50% acetonitrile) in a volatile high-pH elution solution was then added to the columns to elute the peptides as eight different gradient fractions. The fractionated samples were then evaporated using vacuum evaporation and resuspended in 20 μL of 0.1% formic acid. Two microliters of each fraction were loaded for LC-MS/MS analysis.

### Liquid chromatography and Mass spectrometry

The iRT reagent (Biognosys, Switzerland) was added at a ratio of 1:10 v/v to all peptide samples to calibrate the retention time of the extracted peptide peaks. For analysis, 1 μg of the peptide from each sample was loaded into a trap column (75 μm * 2 cm, 3 μm, C18, 100 Å) at a flow rate of 0.55 μL/min and then separated with a reversed-phase analytical column (75 μm * 250 mm, 2 μm, C18, 100 Å). Peptides were eluted with a gradient of 3%–90% buffer B (0.1% formic acid in 80% acetonitrile) for 120 min and then analyzed with an Orbitrap Fusion Lumos Tribrid Mass Spectrometer (Thermo Fisher Scientific, Waltham, MA, USA). 120 Min gradient elution: 0 min, 3% phase B; 0 min-3 min, 8% phase B; 3 min-93 min, 22% phase B; 93 min-113 min, 35% phase B; 113 min-120 min, 90% phase B. The LC settings were the same for both the DDA-MS and DIA-MS modes to maintain a stable retention time.

For the generation of the spectral library (DIA), the eight fractions obtained from the spin column separation were analyzed with mass spectrometry in DDA mode. The MS data were acquired in high-sensitivity mode. A full MS scan was acquired within a 350–1200 m/z range with the resolution set to 120,000. The MS/MS scan was acquired in Orbitrap mode with a resolution of 30,000. The HCD collision energy was set to 30%.

The AGC target was set to 4e5, and the maximum injection time was 50 ms. The individual samples were analyzed in DDA/DIA-MS mode. The variable isolation window of the DIA method with 29 windows was used for DIA acquisition. The full scan was obtained at a resolution of 120,000 with an m/z range from 400 to 1200, and the DIA scan was obtained at a resolution of 30,000. The AGC target was 1e5, and the maximum injection time was 50 ms. The HCD collision energy was set to 35%.

### Mass spectrometry data processing

The Ms data of the rat cohort was performed label-free quantitative comparisons. Base peak chromatograms were inspected visually in Xcalibur Qual Brower version 4.0.27.19(Thermo Fisher Scientific). To generate a spectral library, ten DDA raw files were first searched by Proteome Discoverer (version 2.1; Thermo Scientific) with SEQUEST HT against the Uniprot rat sequence database (April 17, 2021; 36,181 sequences). The iRT sequence was also added to the human database. The search allowed two missed cleavage sites in trypsin digestion. Carbamidomethyl (C) was specified as the fixed modification. Oxidation (M) was specified as the variable modification. The parent ion mass tolerances were set to 10 ppm, and the fragment ion mass tolerance was set to 0.02 Da. The Q value (FDR) cutoff at the precursor and protein levels was 1%. Then, the search results were imported to Spectronaut Pulsar (Biognosys AG, Switzerland) software to generate the spectral library (Bruderer et al., 2015).

The individual acquisition DIA files were imported into Spectronaut Pulsar with default settings. The peptide retention time was calibrated according to the iRT data. Cross-run normalization was performed to calibrate the systematic variance of the LC-MS performance, and local normalization based on local regression was used (Callister et al., 2006). Protein inference was performed using the implemented IDPicker algorithm to generate the protein groups (Zhang et al., 2007). All results were then filtered according to a Q value less than 0.01 (corresponding to an FDR of 1%). The peptide intensity was calculated by summing the peak areas of the respective fragment ions for MS2. The protein intensity was calculated by summing the respective peptide intensity.

The permutation combination comparison method was used to analyze every rat data individually of all timepoints. The methods included two means: comparing two adjacent timepoints and comparing that phase timepoint with the previous phase timepoint (**as shown in Figure 2**). The differential proteins were screened with the following criteria: proteins with at least two unique peptides were allowed; fold change ≥2 or ≤ 0.5; and P < 0.05 by Student’s t-test. Group differences resulting in P < 0.05 were identified as statistically significant. The P-values of group differences were also adjusted by the Benjamini and Hochberg method (Benjamini and Hochberg, 1995). The differential proteins were analyzed by Gene Ontology (GO) based on biological processes(BP), cellular components(CC), and molecular functions(MF) using DAVID (Da Huang et al., 2009), and biological processes from WebGestalt (http://www.webgestalt.org). Protein interaction network analysis was performed using the STRING database (https://string-db.org/cgi/input.pl) and visualized by Cytoscape (V.3.7.1) (Franz et al., 2016) and OmicsBean workbench (http://www.omicsbean.cn).

## Competing interests

The authors declare no competing interests.

## Funding

the National Key Research and Development Program of China (2018YFC0910202, 2016YFC1306300), the Fundamental Research Funds for the Central Universities (2020KJZX002), Beijing Natural Science Foundation (7172076), Beijing cooperative construction project (110651103), Beijing Normal University (11100704), Peking Union Medical College Hospital (2016-2.27).

## Data Availability

The mass spectrometry proteomics data have been deposited to the ProteomeXchange Consortium (http://proteomecentral.proteomexchange.org) via the iProX partner repository (Ma et al., 2019) with the dataset identifier.

## Notes

### Competing Interest Statement

The authors have declared no competing interest.

